# Integrative proteomics and transcriptomics of human T-cells reveals temporal metabolic reprogramming following TCR-induced activation

**DOI:** 10.1101/2023.03.17.532022

**Authors:** Harshi Weerakoon, Ahmed Mohamed, Yide Wong, Bhagya Senadheera, Oscar Haigh, Thomas S. Watkins, Stephen Kazakoff, Pamela Mukhopadhyay, Jason Mulvenna, John J. Miles, Michelle M. Hill, Ailin Lepletier

## Abstract

T-cells are critical components of the adaptive immune system. Upon activation, they acquire effector functions through a complex interplay between mRNA transcripts and proteins, the landscape of which remains to be fully elucidated. In this resource article, we present an integrative temporal proteomic and transcriptomic analysis of primary human CD4^+^ and CD8^+^ T-cells following *ex vivo* activation with anti-CD3/CD28 Dynabeads. Our data reveal a time-dependent dissociation between the T-cell transcriptome and proteome during activation. A transient downregulation of GLUT1, the central glucose transporter in T-cells, marked the onset of reprogramming in both CD4^+^ and CD8^+^ T-cells. At late activation, CD4^+^ T-cells upregulated enzymes associated with degradation of fatty acids while CD8^+^ T-cells preferentially upregulated enzymes in the metabolism of cofactors and vitamins. Surprisingly, we found that activated CD4^+^ and CD8^+^ T-cells became transcriptionally more divergent at the same time their proteome became more similar. In addition to the metabolic reprogramming highlighted in our analysis, this dataset provides a public resource for understanding temporal molecular changes governing the acquisition of effector functions by T-cells.

## Background

T-cells are key players in adaptive immunity and have a major role in the surveillance against pathogens and tumor cells while maintaining unresponsiveness to self-antigens. The metabolic and protein synthesis machinery that shape T-cell responses is controlled by immune activation. Stimulation of the T-cell receptor (TCR) and its costimulatory molecule, CD28, initiates a transcriptional program in naïve T-cells that leads to activation, expansion and differentiation into specialized CD4^+^ “helper” and CD8^+^ “cytotoxic” T-cells (Shah *et al*, 2021; Kumar *et al*, 2018). The duration of TCR signaling is reported as a key factor determining the functional qualities of the T-cells that develop and their commitment to proliferation (Shinzawa *et al*, 2022; Prlic *et al*, 2006; Au-Yeung *et al*, 2014). In this regards, the time of TCR stimulation necessary to launch the proliferative program for naive CD4^+^ T-cells has been shown to be more than required to CD8^+^ T-cells (Kaech & Ahmed, 2001; Iezzi *et al*, 1998).

The extensive reprogramming of activated T-cells reflects substantial remodeling of multiple molecular pathways involved in cellular metabolism and protein synthesis increasingly being comprehended due to recent advancements in the fields of proteomics, transcriptomics, metabolomics and epigenomics (Papale, 2021; Wang *et al*, 2008; Shay & Kang, 2013; Hukelmann *et al*, 2016; Tan *et al*, 2017). Initial genomic and transcriptomic studies laid the baseline for understanding the T-cell reprogramming following TCR stimulation but only captured a partial snapshot of this complex process, also involving several protein-protein interactions and protein phosphorylation (Soskic *et al*, 2022; Shifrut *et al*, 2018). Our knowledge on changes in the proteome of activated T-cells has widened with the application of high throughput proteomic technologies in the field of immunology (Weerakoon *et al*, 2020; Subbannayya *et al*, 2021; Suomi & Elo, 2022; Papale, 2021). Based on proteomics analysis of mouse primary cells, the competitive proliferative advantage of activated CD8^+^ over CD4^+^ T-cells was found to be associated with differences in their intrinsic nutrient transport and biosynthetic capacity (Howden *et al*, 2019).

The caveat of studies based on the traditional single-omics approach is the limitation in providing integrated mRNA-protein data, which offers the opportunity to understand the flow of information that underlies T-cell activation and the acquisition of specialized effector functions. Further, many investigations in the fields of T-cell immunology and biology originate from observations based on animal experimental models and T-cell lines limiting the current knowledge around the immune response of primary human T-cells to TCR stimulation (Mestas & Hughes, 2004; Mak *et al*, 2014). As such, mapping the T-cell activation cycle at multiple molecular levels is crucial to understand the complex mechanisms underpinning this process, as well as identifying conditions where TCR activation is disrupted by negative signals that influence the quality of T-cell responses.

To characterize the main pathways governing the acquisition of effector functions by CD4^+^ and CD8^+^ T-cells, we have generated a temporal proteomic and transcriptomic dataset of human primary T-cell activation. Here, we report an integrative analysis of the dataset that revealed the extent of transcriptome-proteome discordance in activated T-cells, the differentially expressed pathways between CD4^+^ and CD8^+^ T-cells, and the metabolic reprogramming during the different phases of activation. Furthermore, this temporal dataset of human primary T-cell activation provides an important public reference resource for immunology and biomedical research.

## Results

### Uncoupling of T-cell proteome and transcriptome following activation

There has traditionally existed an assumed implication of a proportional relationship between mRNA transcription abundance and protein expression measured from a tissue. However, concomitant analysis of gene and protein expression can frequently fail to provide a correlation between the two domains (Marguerat *et al*, 2012; Jovanovic *et al*, 2015; Payne, 2015; Zhang *et al*, 2014; Saelao *et al*, 2018; Johansson *et al*, 2019). To explore the temporal transcriptomic and proteomic changes mediated by TCR activation in both CD4^+^ and CD8^+^ T-cells, these subsets were purified from the blood of three healthy volunteers and analyzed at different time points following activation with CD3/CD28 Dynabeads; 0 hour (h), 6h, 12h, 24h, 3 days (d), and 7d (Figure 1A), using a parallel workflow to generate both transcriptomic and proteomic datasets. These time points were chosen to represent early (up to 24h) and late phase (3d and 7d) of T-cell activation. Prior to RNA sequencing (RNA-seq) and label-free data-dependent acquisition mass spectrometry-based proteomics (DDA-proteomics), the purity of isolated CD4^+^ and CD8^+^ T-cells was assessed by fluorescence-labeled flow cytometry (FACS) and monoclonal antibodies to be >90% (Supplementary Figure 1A).

**Figure 1.**
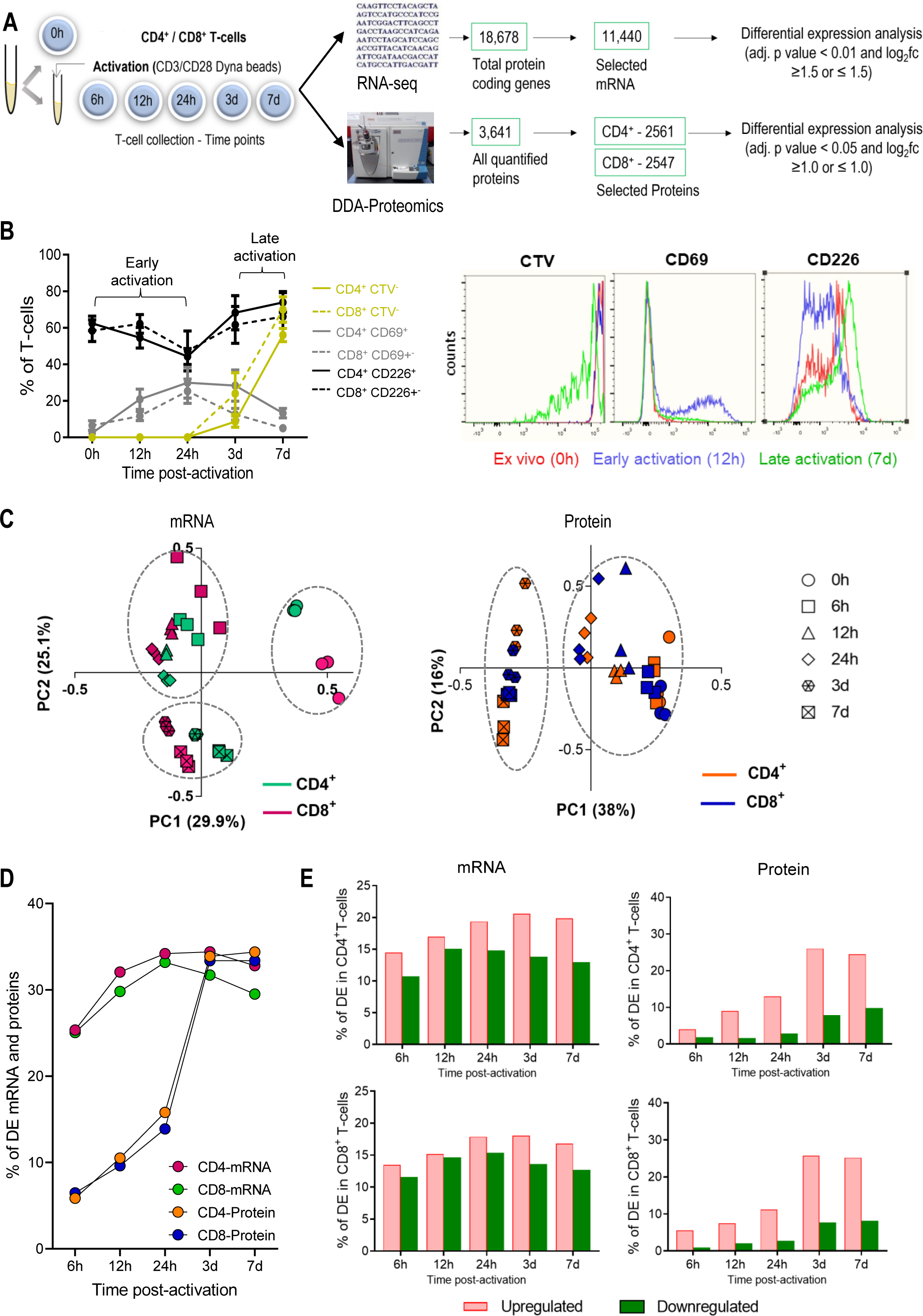
Minor changes to T-cell proteome during early stages of T-cell activation. **A.** Schematic of CD4^+^ and CD8^+^ T-cells isolation, activation, and parallel transcriptomic and proteomic analysis at 6 time points using RNA sequencing (RNA-seq) and data dependent acquisition proteomic analysis (DDA-proteomics), respectively. The number of transcripts/proteins pre-and post-quality control is indicated, along with criteria for differential expression (DE) analysis. **B.** Characterization of stages of T-cell activation through T-cell activation markers, CD69 and CD226, and proliferative T-cells (CTV^-^) by flow cytometry; early activation (CD69^high^CD226^low^CTV^+^) and late activation (CD69^high^CD226^low^CTV^-^). **C.** Principal component analysis shows the relationship of mRNA and protein data from three biological replicates across different time points. **D.** Ratio of mRNA and protein differentially expressed at different time points in relation to their corresponding unstimulated controls (0h = 1). **E.** Bar charts indicate DE genes and proteins in CD4^+^ T-cells and CD8^+^ T-cells as a percentage of total mRNA/ proteins detected in each time point in relation to unstimulated cells. red: up-regulated, green: down-regulated mRNA/ proteins.

A total of 18,678 protein-coding mRNA and 3,531 proteins were identified from a total of 36 samples analyzed (Figure 1A). Quality control measures computed using RNA-SeQC and Maxquant output ensured the suitability of the refined transcriptome and proteome datasets, respectively. Based on similar distribution of mRNA copy numbers (Supplementary Figure 1B), similar total protein intensities (Supplementary Figure. 1C), as well as < 20% missing values for proteins (Supplementary Figure 1D), 11,440 mRNA transcripts (Supplementary Table 1) and ∼2,550 proteins (Supplementary Table 2) were selected for differential expression (DE) analysis.

To define the temporal dimensions of T-cell activation, we assessed cell proliferation (based on cell trace violet (CTV) dilution) alongside cell activation (based on the detection of surface markers for early (CD69) and late (CD226) phases of T-cell activation) (Lepletier *et al*, 2019). In line with previous studies (Obst, 2015), T-cells started to proliferate after 24h of stimulation (Figure 1B). In both CD4^+^ and CD8^+^ T-cells, a pronounced CD69 surface expression was elicited between 6h and 24h, followed by gradual reduction. Conversely, CD226 expression was transiently downregulated in the first 24h (Figure 1B), indicating 24h as the inflection point between the early and late activation phases in our dataset. This was further supported by principal component (PC) analyses, which revealed that mRNA obtained from unstimulated (0h), early-or late-activated T-cells, formed three well defined and distinct clusters (Figure 1C, left). A similar protein clustering pattern was observed between unstimulated and early activated T-cells (Figure 1C, right).

DE analysis was conducted by comparing each activation time point to 0h, revealing the percentage of mRNA or proteins that were differential in each T-cell type over time (Figure 1D and 1E). As expected, the overall changes in the T-cell transcriptome content preceded changes at the protein level. As early as 6h following activation, the expression of ∼25% of the transcriptome was significantly changed, in contrast to only ∼5% of the proteome, in both T-cell subsets. However, during the proliferation phase (late phase of activation), the fraction of DE mRNA transcripts and proteins became almost equal, due to a dramatic increase in proteins but little change in mRNA contents (Figure 1D).

Together, these data reveal a rapid and drastic T-cell transcriptomic response following TCR activation, which converts to a refined proteomic response with prolonged stimulation that coincided with proliferation.

### Proteome and transcriptome rewiring at late stage of T-cell activation

We hypothesized that the signal propagation required for the conversion of mRNA transcripts into proteins would be temporally regulated during T-cell activation. Supporting this idea, a high discrepancy between the mRNA and protein content was observed during CD4^+^ and CD8^+^ T-cell activation (Figure 2A). Only 20% of mRNA transcripts identified in activated T-cells, were quantified at the proteomic level (Figure 2B), likely due to sensitivity limitations of the proteomic technology. In view of this limitation, we focused on the DE transcripts, with the rationale that a significant increase in transcript abundance should increase the detectability of the cognate protein. From 570 DE transcripts identified at 6h, 150 matching proteins were found to be DE during the course of analysis, representing 25 proteins simultaneously modulated at 6h and over 100 proteins modulated at late phase (Figure 2C). This expression pattern was common to both CD4^+^ and CD8^+^ T-cells and indicates a time delay of at least 3 days for a significant proportion of the mRNA transcripts to be translated into proteins in response to T-cell activation. Interestingly, a gradual and consistent increase was observed towards the later time points analyzed. While the correlation between proteins and mRNA simultaneously expressed at 6 hours was poor (r=0.35 and r=0.23 for CD4^+^ and CD8^+^, respectively), a moderate/strong correlation between mRNA and protein groups was observed at 3d (r=0.67 and r=0.73 for CD4^+^ and CD8^+^, respectively) and 7d (r=0.69 and r=0.72 for CD4^+^ and CD8^+^, respectively) (Figure 2D and 2E).

**Figure 2.**
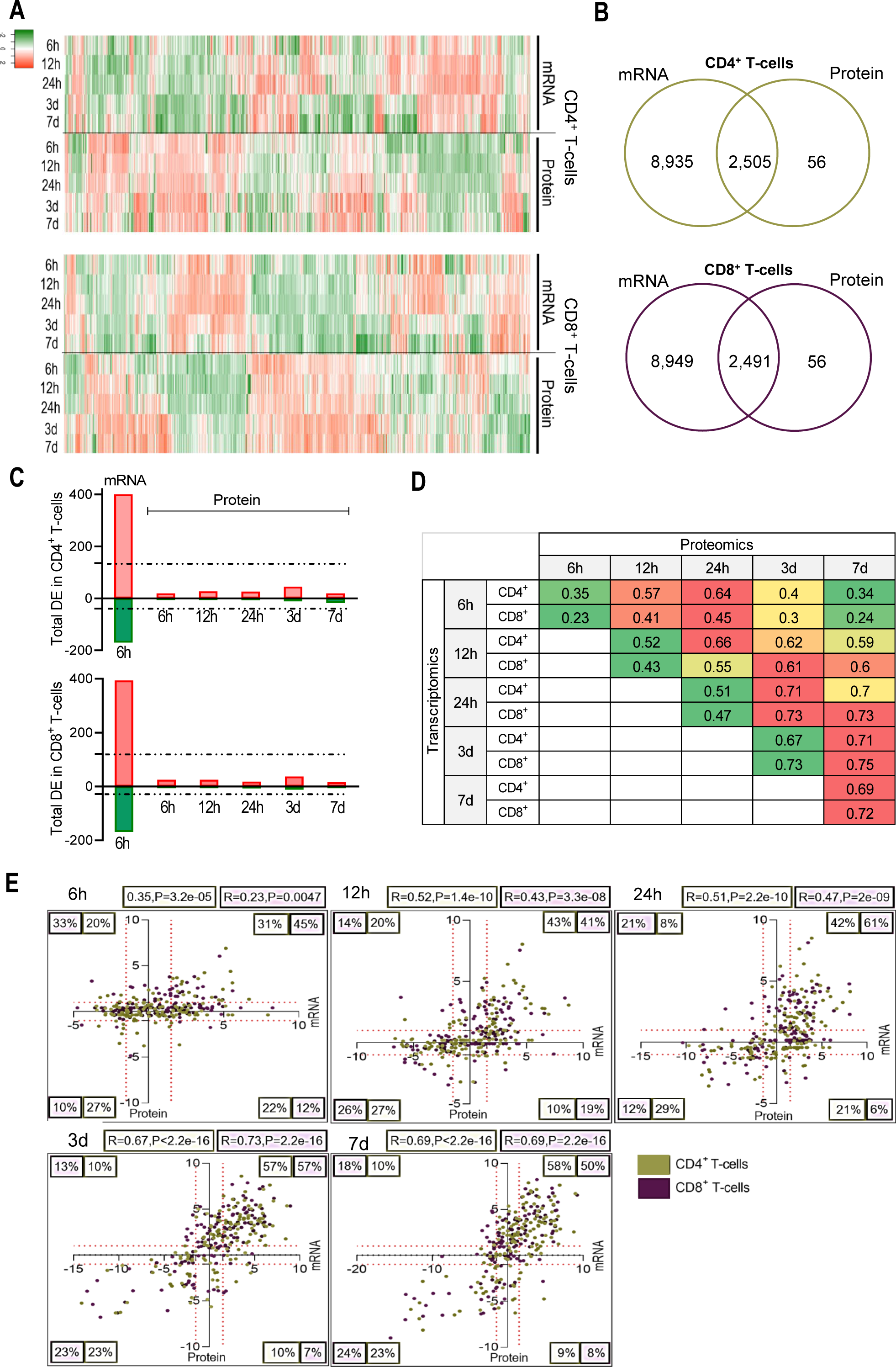
Proteome and transcriptome rewire at late stages of T-cell activation. **A.** Heatmaps show the expression patterns of commonly quantified mRNA transcripts and proteins in CD4^+^ and CD8^+^ T-cells. Red: upregulated, grey: no DE, green: downregulated. **B.** Venn diagrams showing the overlap between quantified mRNA and protein obtained from transcriptomic and proteomic data **C.** DE proteins encoded by mRNA differentially expressed at 6 hours following activation. The number of the proteins regulated at each time point is shown. FDR < 0.05. Dotted line represents the total number of proteins up-(green) or downregulated (red). **D.** Pearson correlation between DE genes and proteins over the entire time course of T-cell activation. Green to red gradient shows low to high correlation values for CD4^+^ and CD8^+^ T-cells at each time point **E.** Scatter graph with four quadrants indicate the distribution and correlation between gene and protein expression changes in both CD4^+^ and CD8^+^ T-cells at different time points following activation. Each region lists the percentage of T-cells falling in each category. mRNA distribution is represented in the “x” axis and protein distribution in the “y” axis. ‘R’ represents Pearson correlation coefficient.

Thus, a lag between the expression of mRNA transcripts and protein synthesis explains a significant part of the transcriptome-proteome discordance observed in both populations of activated T-cells, allowing for a period of temporal regulation and “omic” rewiring.

### TCR activation results in proteomic convergence between CD4^+^ and CD8^+^ T-cells not mirrored at the mRNA level

Comparison of mRNA transcript and protein libraries between activated human CD4^+^ and CD8^+^ T-cells, from a temporal perspective, has not been previously reported. Therefore, we sought to reveal molecular differences between both T-cell subsets by comparing their dynamic transcriptomic and proteomic changes during activation. Among the proteins and transcripts commonly quantified in both subsets, 19% of proteins (487/2,544) and 8% of mRNA transcripts (968/11,440) were found to be DE between CD4^+^ and CD8^+^ T-cells (Supplementary Table 3). Most of the DE proteins were identified at 0h (n=172) (Figure 3A). Over-expressed proteins in CD4^+^ T-cells included the RNA demethylase (ALKBH5, log_2_fc = 3.59), methyl-CpG-binding protein (MBD2, log_2_fc=3.62) and mitochondrial protein (MRPL44, log_2_fc=3.59) (Figure 3B) while over-expressed proteins in CD8^+^ T-cells included the regulator complex proteins (LAMTOR5, log_2_fc=4.57), hexosaminidase subunit beta (HEXB, log_2_fc=3.99) and lysosomal enzyme (AGA, log_2_fc=3.38). The transcription regulator Runt-related transcription factor 3 (RUNX3, log_2_fc=3.75) and distinct profiles of cytotoxic granules (granzymes, GZMM and GZMA), were also among the highest upregulated proteins in CD8^+^ T-cells at 0h (Figure 3B). Interestingly, during activation, the expression of proteins highly DE at 0h became more similar between CD4^+^ and CD8^+^ T-cells (Figure 3C, top graphs), while the expression of their corresponding transcripts did not significantly change (Figure 3C, bottom graphs). Despite reduction in the number of DE proteins, CD8^+^ T-cells maintained proteins associated with cytolysis enriched during the entire time course analyzed (Supplementary Figure 2A).

**Figure 3.**
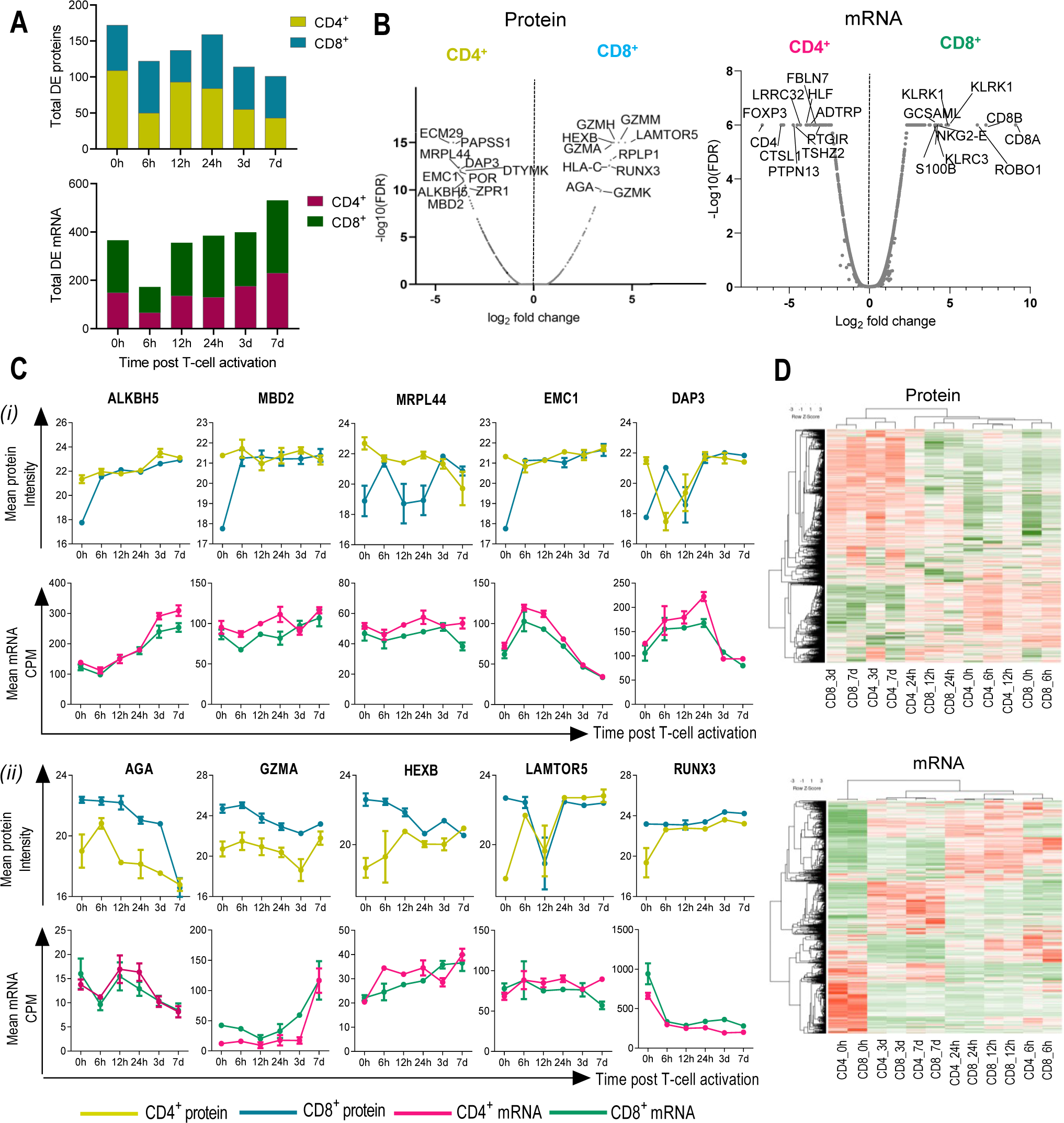
CD4^+^ and CD8^+^ T-cells become more divergent during activation. **A.** Stacked bar graphs showing total number of proteins and mRNA transcripts DE between CD4^+^ and CD8^+^ T-cells at each timepoint. Columns represent transcripts and proteins overexpressed in each T-cell subset. **B.** Volcano plots showing proteins mRNA DE between CD4^+^ and CD8^+^ T-cells at 0 hours. Name of top 10 overexpressed mRNA and proteins in CD8^+^ and CD4^+^ T-cells are indicated. **C.** Expression kinetics of proteins upregulated in CD4^+^ (*i*) and CD8^+^ (*ii*) T-cells and their corresponding mRNA transcripts. Intensities of each time point are shown as mean and the standard error of mean (SEM) (n=3). **D.** Heat map shows the relationship of protein/mRNA expression between CD4^+^ and CD8^+^ T-cells over the time course. mRNA and protein commonly quantified between two T-cell subsets were used. Average linkage and Pearson distance measurement were used in column clustering.

Among the mRNA transcripts overexpressed in CD8^+^ T-cells were the canonical markers CD8A and CD8B, natural killer (NK) cell receptors (KLRK1, KLRC3 and NKG2-E) (Chen *et al*, 2020), and receptors involved in cytolysis (CRTAM and CD160) (Figure 3B, Supplementary Figure 2B). CD4 and FOXP3, the transcriptional regulator required for the development of regulatory T-cell, were among the overexpressed transcripts in CD4^+^ T-cells (Figure 3B, Supplementary Figure 2C). In contrast to proteins, the mRNA content became more distinct between CD4^+^ and CD8^+^ T-cells following activation, and most of the DE mRNA transcripts were identified at 7d (n=530) (Figure 3A). Supporting lower mRNA discrepancy at 0h, unsupervised hierarchical analysis comparing expression profiles between CD4^+^ and CD8^+^ T-cells, revealed that the identified transcripts clustered across both cell subsets, while expressed proteins did not present a clear clustering pattern at 0h and during early activation (Figure 3D).

The significant reduction in the number of DE proteins at late activation indicates the acquisition of similar phenotypic features between CD4^+^ and CD8^+^ T-cells coinciding with their proliferative states, which was not captured at the mRNA level.

### Integrated omics networks reveal temporal changes in the metabolic pathways of CD4^+^ and CD8^+^ T-cells following activation

To elucidate the main cellular pathways supporting CD4^+^ and CD8^+^ T-cell activation, we used a soft clustering tool to divide mRNA transcripts and proteins that significantly changed following TCR/CD28 activation in 12 clusters, according to their kinetics of expression (Figure 4A, Supplementary Figure 3, Supplementary Table 4). Although clusters were formed with a similar number of transcripts and proteins representing each T-cell subset (Figure 4A, Supplementary Figure 3), the overall expression overlap observed between CD4^+^ and CD8^+^ T-cells identified as part of the same kinetics cluster was intermediate for mRNA transcripts (∼50% overlap) and poor for proteins (∼25% overlap) (Figure 4B).

**Figure 4.**
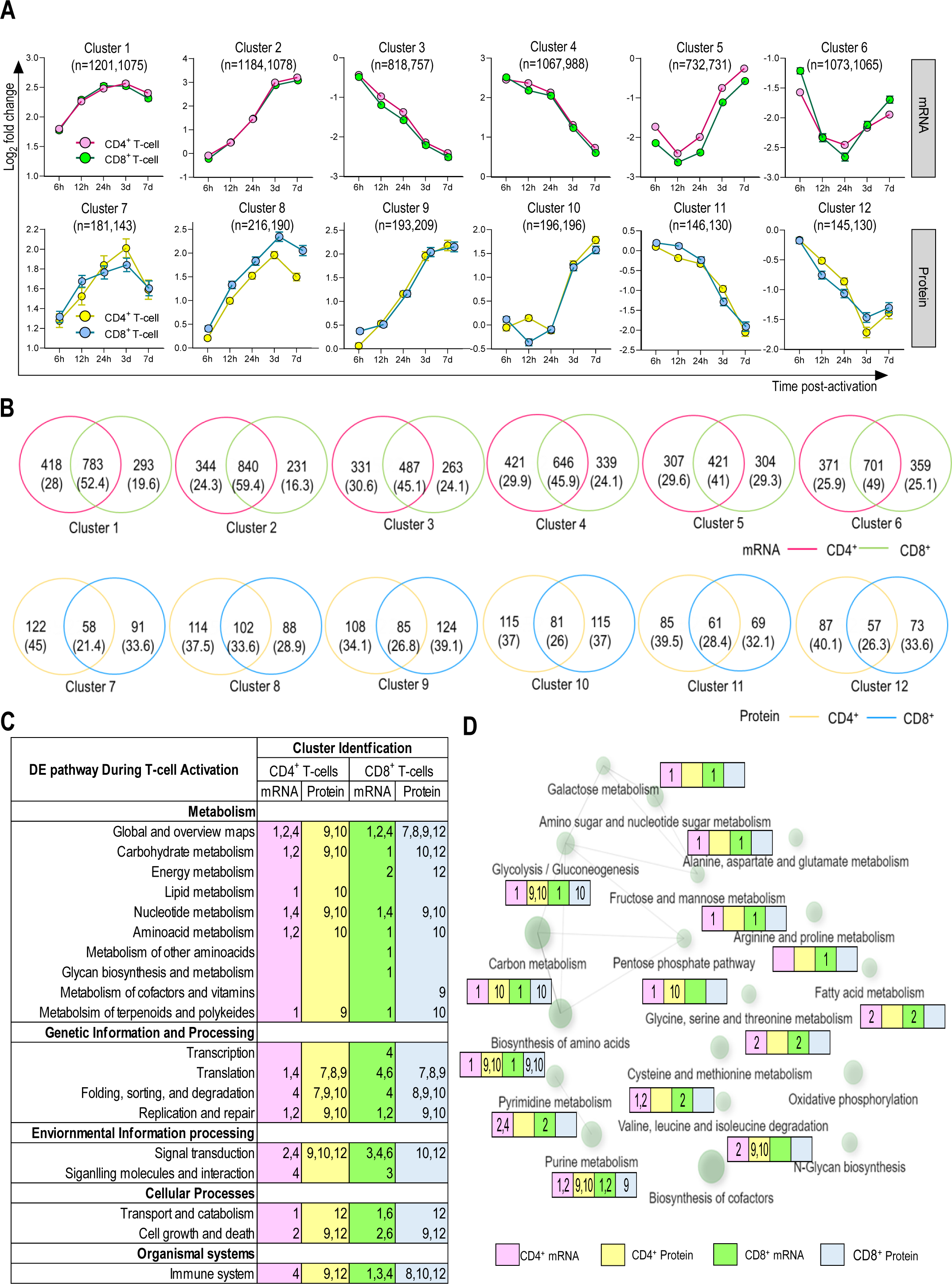
Main pathway altered during T-cell activation is associated with metabolism. **A.** Co-expression clusters of transcriptome and proteome data from CD4^+^ and CD8^+^ T-cells. DE mRNA transcripts of each dataset were clustered using mFuzz soft clustering (R package). mRNA or protein intensities at each time point are shown as mean and the standard error of mean (SEM). Number of mRNA/proteins included in each cluster are indicated for CD4^+^ and CD8^+^ T-cells, respectively. mRNA transcripts with log2fc > 1.5 or < −1.5 and proteins with log_2_fc ≥ 1.0 or ≤ −1.0 were considered as DE. **B.** Venn diagrams showing the overlap of mRNA and protein identified in each cluster between activated CD4^+^ and CD8^+^ T-cells. **C.** Enriched KEGG pathways (FDR < 0.05) for co-expression clusters (defined in Figure 4A) of CD4^+^ and CD8^+^ T-cell. **D.** Molecular interaction, reaction and relation network showing the relationship of the top first 20 enriched KEGG pathways categorized under ‘metabolism’ (FDR < 0.05). The network was generated using all DE mRNA transcripts and proteins. The size of each node directly correlates with the number of genes included. Edges represents sharing of 20% or more genes between two nodes while the thickness of the edge directly correlates with the number of overlapping genes. The number in each coloured box indicates the co-expression clusters in Figure 4A from where the corresponding mRNA transcript/ protein was enriched. In pink CD4^+^ mRNA, yellow CD4^+^ protein, green CD8^+^ mRNA, blue CD8^+^ protein.

To map the main biological changes underpinning T-cell activation, we next conducted a functional pathway enrichment analysis using Kyoto Encyclopedia of Genes and Genomes (KEGG). Pathways categorized under metabolism, genetic information processing, environmental information processing, cellular processes, and organismal systems were selected for comparative analysis between activated CD4^+^ and CD8^+^ T-cells (Figure 4C). As expected, major changes in metabolic pathways were detected in activated T-cells at both protein and transcripts levels. Differences in both mRNA transcripts and proteins governing glycolysis/gluconeogenesis, carbon metabolism, and biosynthesis of amino acids, were observed following TCR activation. However, changes in the metabolism of essential (methionine and threonine) and non-essential (alanine, aspartate, glutamate, glycine, serine, and cysteine) amino acids were only captured by transcriptomics in both cell subsets, while DE transcripts associated with arginine and proline metabolism were exclusively detected in CD8^+^ T-cells (Figure 4D). Following activation, the expression of proteins in the lipid metabolism pathway mostly represented by enzymes related with degradation of fatty acids, such as ACAT2, ACSL4, ACADVL and HADH, exponentially upregulated in CD4^+^ T-cells only (cluster 10) (Figure 4A and 4C, Supplementary Table 5). In parallel, proteins associated with energy metabolism and metabolism of cofactors and vitamins were DE solely in CD8^+^ T-cells (clusters 9 and 12, respectively) (Figure 4A and 4C).

As the transport of nutrients from the surrounding environment is a crucial factor in modulating the molecular mechanisms that lead to activation, we mapped the kinetics of the glucose and amino acid transporters to identify T-cell reprogramming following TCR/CD28 activation. Corroborating findings from previous studies (Bevilacqua *et al*, 2022), a number of amino acid receptors involved in glutamine uptake, including the transporters SLC1A5, SLC7A5, and SLC3A2, showed upregulation of their corresponding mRNA and proteins as early as 6h (Figure 5, Supplementary Figure 4A). Interestingly, upregulated transcripts for these transporters reduced to unstimulated T-cell levels after 12h, while their corresponding proteins exponentially increased during the activation time course in both CD4^+^ and CD8^+^ T-cells. Similar expression kinetics was observed for transcripts and proteins representing enzymes in the glutaminolysis pathway: GLS, which convert glutamine into TCA (tricarboxylic acid) cycle metabolites, pyruvate producer ME2, and pyruvate metabolizer LDH (Figure 5).

**Figure 5.**
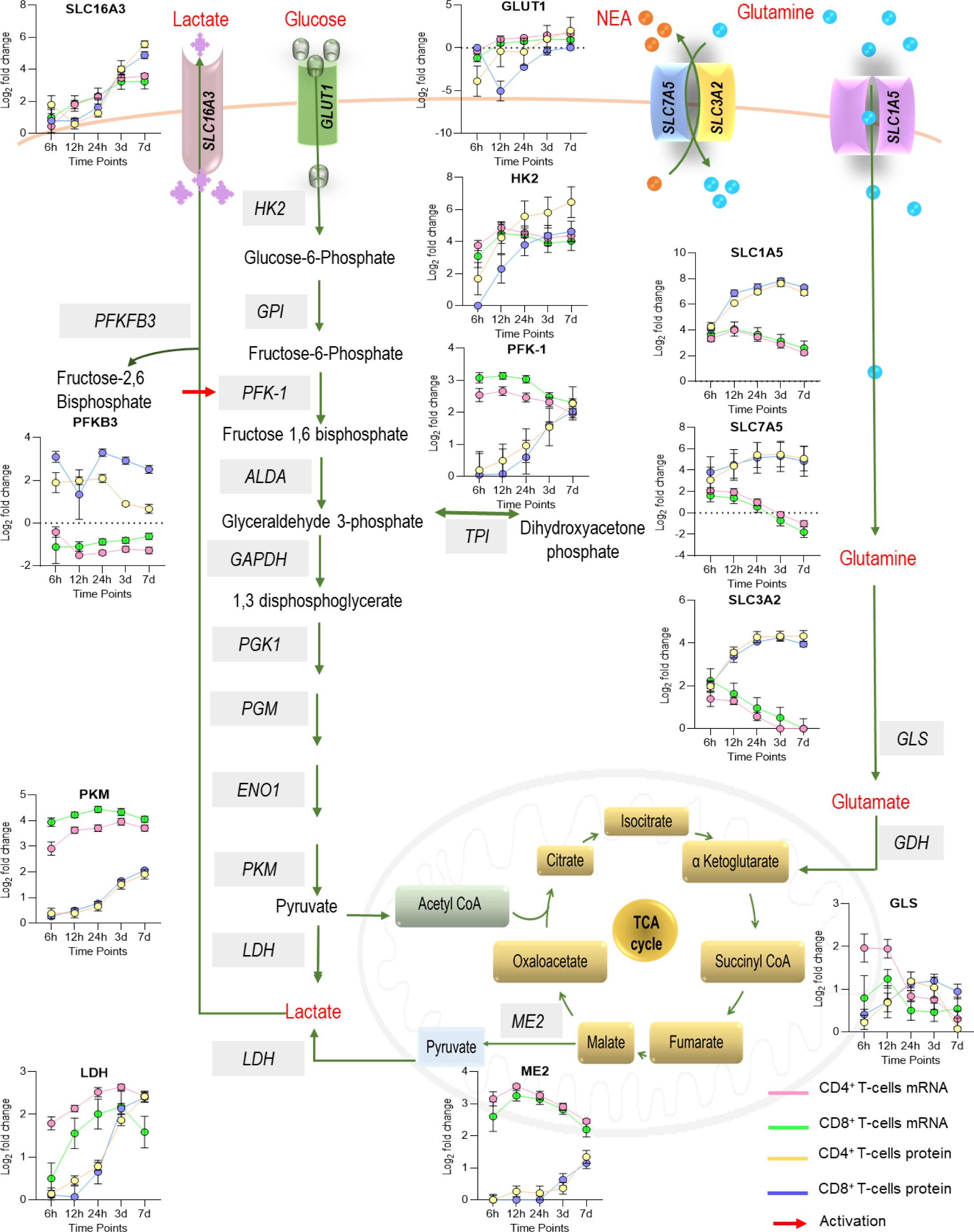
Re-wiring of aerobic glycolysis and glutaminolysis detected by multi-omic analysis during T-cell activation. Besides glucose, T-cells utilize glutamine through the glutaminolysis pathway to produce energy during the activation. Line graphs show the dynamic protein and gene expression patterns of the main glucose and glutamine transporters and the rate-limiting/ key enzymes of aerobic glycolysis and glutaminolysis during CD4^+^ (orange color) and CD8^+^ (blue color) T-cell activation (0 – 24 hours) and proliferation (3^rd^ to 7^th^ day). Expressed proteins are named as follow: GLUT-1 (SLC2A1) - the main glucose transporter in T-cells, SLC7A5, SLC3A2 and SLC1A5 - glutamine transporters, SLC16A3 –lactate transporter, HK2, PFKP and PKM – the rate-limiting enzymes of glycolysis, PFKFB3 – a key allosteric activator of glycolysis, LDH – the enzyme which converts pyruvate to lactate, GLS – the enzyme which converts glutamine to glutamate and ME2 – the enzyme which converts malate to pyruvate in the mitochondrial matrix.

As T-cells are known to perform aerobic glycolysis to fulfill the bioenergetic demands of activation (Almeida *et al*, 2016; Chapman *et al*, 2020), we further explored our datasets to identify DE mRNA transcripts and proteins in the glycolysis pathway. Surprisingly, the observed increase in glutamine transport and metabolism was paralleled by a transitory downregulation of protein measures for the main glucose transporter expressed by T-cells, GLUT1 (SLC2A1), between 6h (CD4^+^ T-cell) and 12h (CD8^+^-T cell) following initial activation (Figure 5, Supplementary Figure 4B). The results obtained from the high throughput proteomics data were validated by flow cytometry (FACS) using a fluorescent labeled monoclonal antibody, demonstrating downregulation of GLUT1 during the first 12h in both CD4^+^ and CD8^+^ T-cells (Supplementary Figure 4B). Strikingly, in comparison to proteomics, the FACS data revealed a much higher increase in GLUT1 expression after 24h, maybe reflecting the difference in surface expression captured by FACS versus total GLUT1 captured by proteomics. The decrease in GLUT1 expression was paralleled by transitory downregulation of PFKB3 (a key allosteric activator of glycolysis) protein in CD8^+^ T-cells only, whilst PFKB3 mRNA transcript remained downregulated during the entire time course analyzed (Figure 5, Supplementary Figure 4C and 4D). Protein expression of SLC16A3, a high affinity transporter capable of exporting lactate and pyruvate in response to the glycolytic influx, transiently dropped at 12h in CD4^+^ T-cells only (Figure 5). In both CD4^+^ and CD8^+^ T-cells, expression of the rate-limiting enzymes of glycolysis, HK2, PKM, and PFK1 increased in the late phase of activation, while their mRNA transcripts were found to be increased as early as 6h and remained elevated at later time points (Figure 5).

Altogether, our data reveal a transient disconnection between the aerobic glycolysis and glutaminolysis pathways during T-cell activation, differently captured across CD4^+^ and CD8^+^ T-cells. Such a finding was only possible due to multi-omic analysis and would be overlooked using the single-omics analysis of the T-cell transcriptome.

## Discussion

Here we report the first temporal transcriptomic and proteomic dataset of primary human T-cell subsets as a reference for probing the molecular events underpinning T-cell activation. Interrogation of these integrated datasets provided novel insights on the molecular reprogramming kinetics of T-cell activation.

Our data indicate a high level of temporal discordance between mRNA transcription and protein expression in T-cells following TCR activation. Interestingly, by the late phase of activation, we observed concordance between the reprogrammed transcriptome and proteome resulting in proliferation. The correlation between mRNA transcript and protein expression can vary according to the cell type and its functional status and a complex discordance in the relationship have been recognized in multiple studies (Marguerat *et al*, 2012; Jovanovic *et al*, 2015; Payne, 2015; Zhang *et al*, 2014; Saelao *et al*, 2018; Johansson *et al*, 2019). A quantitative proteome and transcriptome mapping of paired healthy human tissues from the Human Protein Atlas project revealed that hundreds of proteins could not be detected for highly expressed mRNA and strong differences were observed between mRNA transcripts and protein quantities within and across tissues (Wang *et al*, 2019). This discordance was partly associated with post-translational modifications of proteins induced by external environmental signals, such as the metabolic flux (Buccitelli & Selbach, 2020). We speculate that the discordance observed in this study may be due to insufficient ribosome number/activity to process the rapid transcriptional activation following TCR engagement. This is supported by the observed increase in translation pathway proteins during early activation phase (Figure 4C). While changes in other mRNA silencing mechanisms such as RNA decay (Gratacós & Brewer, 2013), degradation by RNases (Ramanathan *et al*, 2016) or sequestration to stress granules (Mikulits *et al*, 2000; Hoefig *et al*, 2021) were not identified in the pathway analyses, we have not specifically evaluated these mechanisms and therefore cannot definitively exclude them at this stage.

T-cell subsets have differential requirements for energy and biosynthetic precursors during activation. Therefore, differential programming of key metabolic processes such as glycolysis, fatty acid and mitochondrial metabolism can direct T-cell to particular effector functions (Jones *et al*, 2017). As an example, a metabolic shift from oxidative to glycolytic pathways upon engagement of TCR ensures long-term T-cell survival and fuels fast energy supply for biosynthesis and replication (Pearce *et al*, 2013). Our integrated temporal transcriptomics and proteomics design comparing CD4^+^ and CD8^+^ T-cells from the same donors uncovered previously uncharacterized selective transcriptional and translational metabolic reprogramming. Following TCR activation, expression of key enzymes in carbohydrate and energy pathway decrease in CD8^+^ T-cells. At the same time, CD4^+^ T-cells engages enzymes associated with fatty acid degradation to generate acetyl-CoA for the TCA cycle (Figure 4) and upregulate LAMTOR5 (Figure 3C) to activate the mTOR pathway, increasing glycolysis and oxidative phosphorylation required for cytokine production. These data show that CD4^+^ T-cells have higher mitochondrial respiratory capacities compared to CD8^+^ T-cells. We reveal that among the top 10 proteins upregulated in unstimulated CD4^+^ T-cells were the death-associated protein 3 (Dap-3), and a subunit of the endoplasmic reticulum (ER) membrane protein complex (EMC1), which have not been previously characterized in T-cells and may play a role in CD4^+^ T-cell biology. Interestingly, we found that the protein content of CD4^+^ T-cells became gradually more similar to CD8^+^ T-cells over time, including the acquisition of cytotoxic functions by CD4^+^ T-cells, as characterized by increased levels of GZMA and GZMM. While CD4^+^ T-cells with cytotoxic activity able to secrete GZMB and perforin have been observed in various immune responses (reviewed in (Takeuchi & Saito, 2017)), a role for GZMA- and GZMM-expressing CD4^+^ T-cells is less known. It is possible that the reduction observed in the number of DE proteins between both T-cell subsets is associated with the activation method employed in this study. Even though anti-CD3/CD28 Dynabeads generate a more physiologically relevant activation over traditional stimulation methods, such as mitogenic lectins (Trickett & Kwan, 2003), it is likely that the bulk (polyclonal) response subsequent to this type of activation is leading to similar developmental program in both T-cell subsets, as opposed to conditions where activation is achieved directly through antigen-specific interactions. Supporting this concept, naturally recognized peptides have been shown to produce a different metabolic signature to anti-CD3/CD28 stimulation, associated with their TCR binding affinity (Jones *et al*, 2017).

We report a previously uncharacterized transitory downregulation of GLUT1 during early activation. Although the mechanism controlling transitory GLUT1 downregulation is not demonstrated in this study, GLUT1 surface trafficking is known to be regulated through the co-stimulatory receptor CD28, and tight regulation of this transporter is suggested to be imperative for normal T-cell activation (Jacobs *et al*, 2008). Despite the decreased expression of the main glucose transporter, glycolysis in activated T-cells has been shown to occur independently of the glucose influx (Menk et al, 2018). Both lipids and amino acids can be converted into various intermediates of glycolysis and the TCA cycle, allowing them to slip into the cellular respiration pathway through a multitude of side doors including glutaminolysis, the process where glutamine (the most abundant amino acid found in the human body) is converted into mitochondrial TCA cycle intermediates (Chen & Chen, 2022; Song et al, 2020). Correlating with the absolute requirement for glutamine to supply carbon and nitrogen to fuel energy necessary for the synthesis of macromolecules in proliferating T-cells (Carr et al, 2010), our data show an exponential increase in glutamine transporters and glutaminolysis enzymes in activated T-cells. Additionally, we confirm upregulation of the glutamine transporters, SLC1A5, SLC7A5, SLC3A2, alongside glutamate synthase (GLS), the key enzyme able to provide glutamine-derived carbons to the TCA cycle (reviewed in Ren et al, 2017). These findings indicate that glutamine may play an important role in fulfilling the early metabolic requirement unleashed by TCR activation when intracellular levels of glucose are likely to be low. Supporting the idea of a crosstalk between GLUT1 and glutamine transporters in activated T-cells, similarly to glucose uptake, transport of glutamine into T-cells is dependent on CD28 co-stimulation (Jacobs *et al*, 2008; Carr et al, 2010).

The increase in glycolysis and mitochondrial respiration may lead to the accumulation of pyruvate in activated T-cells. Accordingly, we evidence upregulation of the mitochondrial malic enzyme (ME2) during late activation, showing that most of the malate originated from the TCA cycle is likely to be converted to pyruvate rather than oxaloacetate. As pyruvate is rapidly converted into lactic acid in the cell cytoplasm, rather than oxidized in the mitochondrial TCA cycle, rise in intracellular lactate level will cause premature cell death (Chapman *et al*, 2020; Madden & Rathmell, 2021). Our data demonstrate a compensatory mechanism to protect the proliferating CD4^+^ and CD8^+^ T-cells from acidosis, mediated by upregulation of the lactate transporter SLC16A3, which allows for efflux of lactate. Interestingly, extracellular lactate correlates with T-cell proliferation (Grist *et al*, 2018) and additional research is important to investigate possibilities of lactate recycling for the production of energy, as shown for other human cell types (Leverve & Mustafa, 2002).

In summary, we report the first matched temporal transcriptomic and proteomic dataset from CD4^+^ and CD8^+^ T-cells upon TCR activation. Integrated analysis of transcript and protein expression changes across early and late phases, revealing the complexity and differences of CD4^+^ and CD8^+^ T-cells metabolic reprogramming in response to the same generic stimuli. While the current datasets are limited by the depth of proteomic coverage and use of a single stimulus, they nevertheless provide a novel resource for human immunology research on T-cell activation in health and disease.

## ACKNOWLEDGMENTS

We thank the volunteers for donating blood samples for this study. HW was supported by an International Postgraduate Research Scholarship, The University of Queensland, Brisbane, Australia and PhD Top-up Scholarship, QIMR Berghofer Medical Research Institute, Brisbane, Australia. JJM is supported by a NHMRC CDF Level 2 Fellowship (1131732). The authors thank the expertise of staff from the QIMR Berghofer Analytical facility and Dr Nic Waddell for her support in the generation of RNAseq data.

## Material & Methods

### Human CD4^+^ and CD8^+^ T-cell isolation and in vitro activation

Human PBMCs isolated from three healthy young adult volunteer blood donors (age 30-35 years, 2 females, 1 male) were further purified using human pan T-cell isolation kit and magnetic activated cell sorting (MACS) (Miltenyi Biotech, Germany) to isolate unlabeled CD3^+^ T-cells. Approximately 30% of total CD3^+^ T-cells were purified using human CD4^+^ T-cell isolation kit while the rest were sorted with human CD8^+^ T-cell isolation kit (Miltenyi Biotech, Germany) using MACS to obtain untouched CD4^+^ and CD8^+^ T-cells, respectively. The sorted cell populations had purity of over 90%, as assessed by FACS. From each sample, 10^6^ cells were aliquoted for *ex vivo* proteomics and transcriptomics, respectively. The remainder (∼ 7.5 x 10^6^ cells each) were harvested in Roswell Park Memorial Institute (RPMI) 1640 medium supplemented with 10% Foetal calf serum (Gibco, USA) and 50 units/ml penicillin and 50μg/ml streptomycin (Gibco, USA) and activated with ‘Dynabeads human T-cell activator CD3/CD28 (Thermo Fisher Scientific, USA) at the bead: cell ratio of 1:1 as per the manufacturer’s instructions. CD8^+^ T-cell cultures were supplemented with 120 IU/ml human recombinant IL-2 (Sigma Aldrich, USA). T-cells were aliquoted into five samples with 1.5 x 10^6^ cells in each condition to obtain cells at five different time points (6 hours, 12 hours, 24h, 3d and 7d) and incubated at 37^0^C in a humidified, 5% CO_2_ incubator. The culture medium was changed on day 4 of post activation. At each time point, aliquots of activated T-cells were collected washed three times with phosphate buffered saline (PBS), and stored for batch proteomics/transcriptomics processing. T-cells for proteomics were stored at −80^0^C. For transcriptomics, cells were lysed in 400 μl of cold TRIzol before storage at −80^0^C until RNA processing.

### Monitoring the *in vitro* T-cell activation process

T-cell activation and expansion were monitored using T-cell activation markers and a T-cell proliferation assay. In parallel to the main experiment, a sample of CellTraceTM Violet (CTV) (Thermo Fisher Scientific, USA) stained T-cells (CD4^+^ and CD8^+^) from each donor was performed as per the protocol given by the manufacturer. At each time point a sample of cells was stained with CD69-PE-cy7 (BD biosciences, USA) and CD226-FITC (BD Biosciences, USA) along with CD3-APCe780 (eBioscience, USA), CD4-BV711 (BD Bioscience, USA) and CD8-Percp cy5.5 (Bioledgend, USA). Samples were analyzed using FACS to determine dynamic expression changes. The percentage of CTV-cells were analyzed to determine the T-cell proliferation rate at different time points.

### RNA extraction, mRNA library generation and next generation sequencing (NGS)

Total RNA was extracted using TRIzol (Thermo Fisher Scietific, USA) phase separation, following the protocol given by the manufacturer. Ultrapure glycogen (Thermo Fisher Scientific, USA) was used to precipitate total RNA. The quality and quantity of the extracted RNA were analyzed using qubit fluorometer (Invitrogen, Thermo Fisher Scientific, USA) and Agilent 2100 bioanalyzer (Agilent technologies, USA), respectively. and RIN score of over 8.00 was confirmed for all the samples (n = 36). In each sample, 300 ng of total RNA was aliquoted and mRNA libraries were prepared using TruSeq stranded mRNA library preparation kit (Illumina, USA). After quality and quantity assessment of the generated libraries, next-generation sequencing (NGS) was performed using NextSeq 500/550 high output v2 kit (150 cycles) (Illumina, USA) to obtain 800 x 10^6^ paired reads per pool (50 x 10^6^ paired reads per sample). Library generation and NGS were performed at the analytical facility, QIMRB.

### Proteomic sample preparation and LC-MS/MS data acquisition

T-cells (1x10^6^) were lysed in 2% SDS in 100 mM TEAB in the presence of protease inhibitor cocktail. After assessing the protein quantity using pierce BCA protein quantification kit (Thermo Fisher Scientific, USA), ∼ 20 μg from each cell lysate was separated and 200 ng of ovalbumin was added as the internal standard. These samples were reduced, alkylated, and digested using trypsin following the method samples as previously described (Weerakoon *et al*, 2020) to obtain the peptides. After desalting, peptides were quantified using microBCA (Thermo Fisher scientific, USA) protein assay to aliquot 1 μg from each sample for MS analysis and were resuspended in MS grade water with 2% acetonitrile, 0.1% formic acid (v/v) to obtain the final volume of 10 μl. These samples were injected to Protecol C18 trap column in Prominence Nano (Shimadzu, Japan) LC system to separate the ions in a Protecol C18 (200Å, 3 μm particle size, 150 mm x 150 μm) column at a flow rate of 1 μl/min over 180 min linear gradient. Solvent A (0.1% formic acid) and solvent B (100% acetonitrile and 0.1% formic acid) were used for the mobile phase. Peptides were eluted in three consecutive linear gradient: 5–10% solvent B over 5 minutes, 10–27% solvent B over 147 minutes and 27–40% solvent B over 10 minutes. Finally, the column was cleaned using 40% to 95% solvent B for 10 minutes. Chromeleon software (version 6.8, Dionex) embedded in Xcalibur software (version 3.0.63, Thermo Fisher Scientific) was used in the nano LC system. Peptides ionized by the nano spray (Thermo Fisher Scientific, USA) ion source (ion spray voltage - 1.75V, heating temperature 285 °C) were analyzed using a Velos Pro Orbitrap mass spectrometer (Thermo Fisher Scientific, USA). In DDA-MS, the MS was controlled and operated in the “top speed” mode using the Xcalibur software to obtain MS1 and MS2 spectral data for peptide ions with charge status between +2 to +4 at 1.96 second window time.

### Transcriptomic data analysis and identification of differentially expressed genes

Sequence reads were trimmed for adapter sequences using Cutadapt (version 1.11) (Martin, 2011) and aligned using STAR (version 2.5.2a) (Dobin *et al*, 2013). to the GRCh37 assembly with the gene, transcript, and exon features of Ensembl (release 89) gene model. Quality control metrics were computed using RNA-SeQC (version 1.1.8) (Deluca *et al*, 2012), while gene expression was estimated using RSEM (version 1.2.30) (Li & Dewey, 2011). Both counts per million (CPM) and trimmed mean of M-values (TMM) methods were used to normalize the gene expression data and differential expression analysis was carried out using edgeR (R package) (Robinson *et al*, 2010). mRNA with log2fc > 1.5 or < −1.5 at adj. p value of < 0.01 were considered as differentially expressed (DE), up- and down-regulated genes, respectively.

### Proteomic data analysis and identification of differentially expressed proteins

After inspecting the quality of generated DDA-MS data using RawMeat (Vast Scientific), raw files were analyzed using MaxQuant software (Cox & Mann, 2008) against UniProt reviewed human proteome database containing 20,242 entries (downloaded on 25th October 2017), UniProt chicken ovalbumin (UniProt ID – P01012) fasta file and the list of common MS contaminants included in the software. maxLFQ (Cox *et al*, 2014) was used to obtain normalized protein intensity data. Peptides and proteins identified at 1% FDR, were further filtered to identify the proteins with single UniProt protein accessions, detected with at least 2 unique or razor peptides, and m-score value with over 5. Protein intensities (log2 transformed) of activated T-cells of different time points were compared with that of unstimulated samples (baseline) using multiple t-tests with false discovery determination by two-stage linear step-up procedure of Benjamini, Krieger and Yekutieli (q value) (Benjamini *et al*, 2006) to obtain statistical significance and their log_2_fc. Proteins with log_2_fc ≥ 1.0 or ≤ −1.0 at q value of ≤ 0.05 were considered as statistically significant up- and down-regulated genes, respectively.

### Correlation analysis, clustering and gene enrichment analysis of differentially expressed mRNA and protein

Commonly quantified mRNA and protein were selected by mapping uniProt IDs of proteomic data with the Ensemble IDs using UniProt Retrieve/ID mapping. Using log_2_fc values Pearson correlation between DE mRNA transcripts and their corresponding protein was calculated for different time points. Correlation co-efficient values ±(1.00 – 0.70) was considered as a strong correlation while ± (0.69 – 0.40) and ± (0.39 – 0.10) were taken as moderate and weak respectively (Schober *et al*, 2018). To identify the co-expression clusters over the course of activation, mRNA or protein significant in at least one time point were filtered and clustered using the Mfuzz soft clustering method (R package) (Kumar & E Futschik, 2007). KEGG pathways (Kanehisa & Goto, 2000) enriched (FDR ≤ 0.05) by the genes represented by each cluster were identified using String: functional protein network analysis version 11.5 (Szklarczyk *et al*, 2019) and ShinyGO 0.76 (Ge *et al*, 2020). Word clouds were generated using the online tool (https://www.wordclouds.com, accessed December, 2021). Bioinformatics analysis and graph generation were done using R Studio (RStudio Team (2020). RStudio: Integrated Development for R. RStudio, PBC, Boston, MA URL http://www.rstudio.com/.) and GraphPad Prism (version 9.2.0 for Windows, GraphPad Software, San Diego, California USA).

## Data and code availability

The mass spectrometry proteomics data have been deposited on the ProteomeXchange Consortium via the PRIDE partner repository with the dataset identifier PXD038810. The RNAseq raw sequence data are not publicly available because participants did not give consent for the data to be publicly released. The RNAseq genecount data is given in Supplementary Table 6.

## Notes

http://proteomecentral.proteomexchange.org/cgi/GetDataset?ID=PXD038810

### Competing Interest Statement

The authors have declared no competing interest.

