## Supplemental Figures for "Integrative proteomics and transcriptomics of human T-cells reveals temporal metabolic reprogramming following TCR-induced activation"

### Supplementary Figure 1

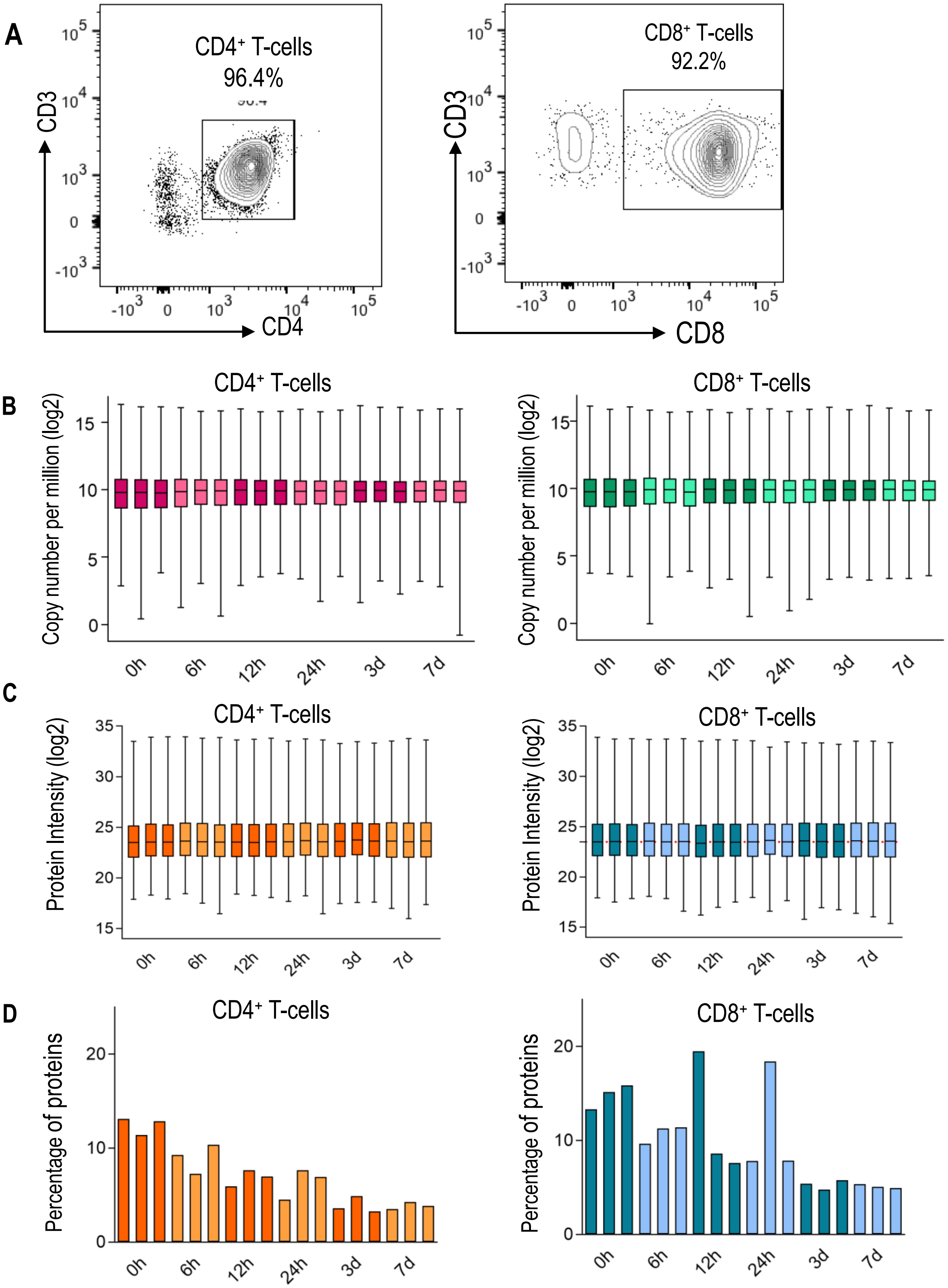


A. Scatter plots obtained from FACS analysis evidencing the purity of sorted $CD4^+$ and $CD8^+$ T-cells

B. Box and Whisker plots showing the the mRNA copy number distribution in each sample (box plots are median, 2nd and 3rd quartiles, whiskers are 95% CI)

C. Distribution of protein intensity data in different $CD4^+$ and $CD8^+$ T-cell samples (box plots are median, 2nd and 3rd quartiles, whiskers are 95% CI)

D. Percentage of proteins with missing intensity values in different $CD4^+$ and $CD8^+$ T-cell samples

### Supplementary Figure 2

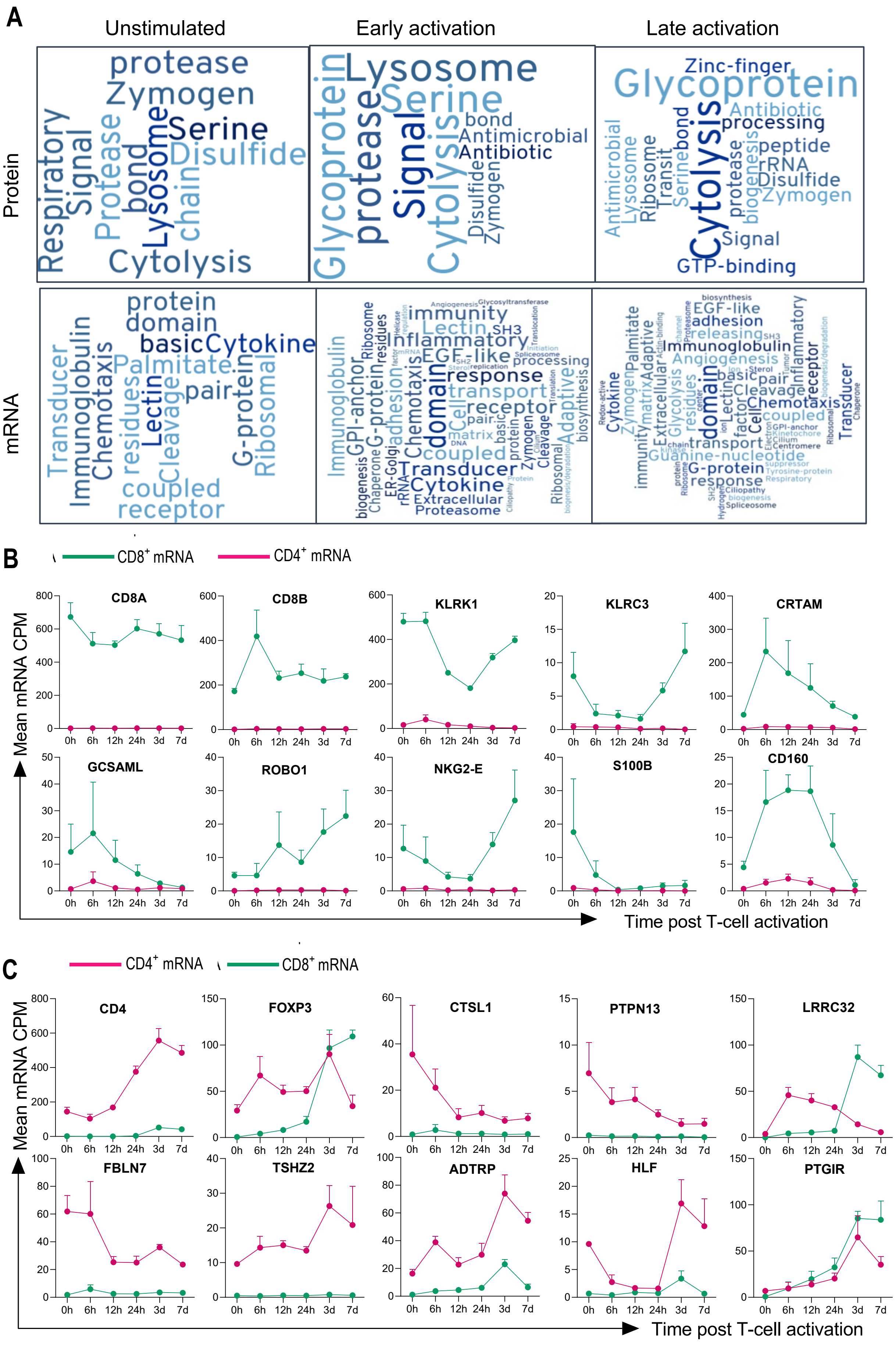

A. Word clouds showing the significant annotated UniProt key words revealed in functional enrichment analysis (FDR < 0.05) of differentially expressed mRNA and protein in $CD8^+$ versus $CD4^+$ T-cells. Early activation word cloud represents annotated UniProt key words identified for 6h, 12h and 24h while late activation represents annotated UniProt key words identified for 3d and 7d.

B. Expression kinetics of the transcripts upregulated in $CD4^+$ T-cells at 0h. mRNA expression values of each activation time point are given as mean and the standard mean of error (SEM).

C. Expression kinetics of the transcripts upregulated in $CD8^+$ T-cells at 0h. mRNA expression values of each activation time point are given as mean and SEM.

### Supplementary Figure 3

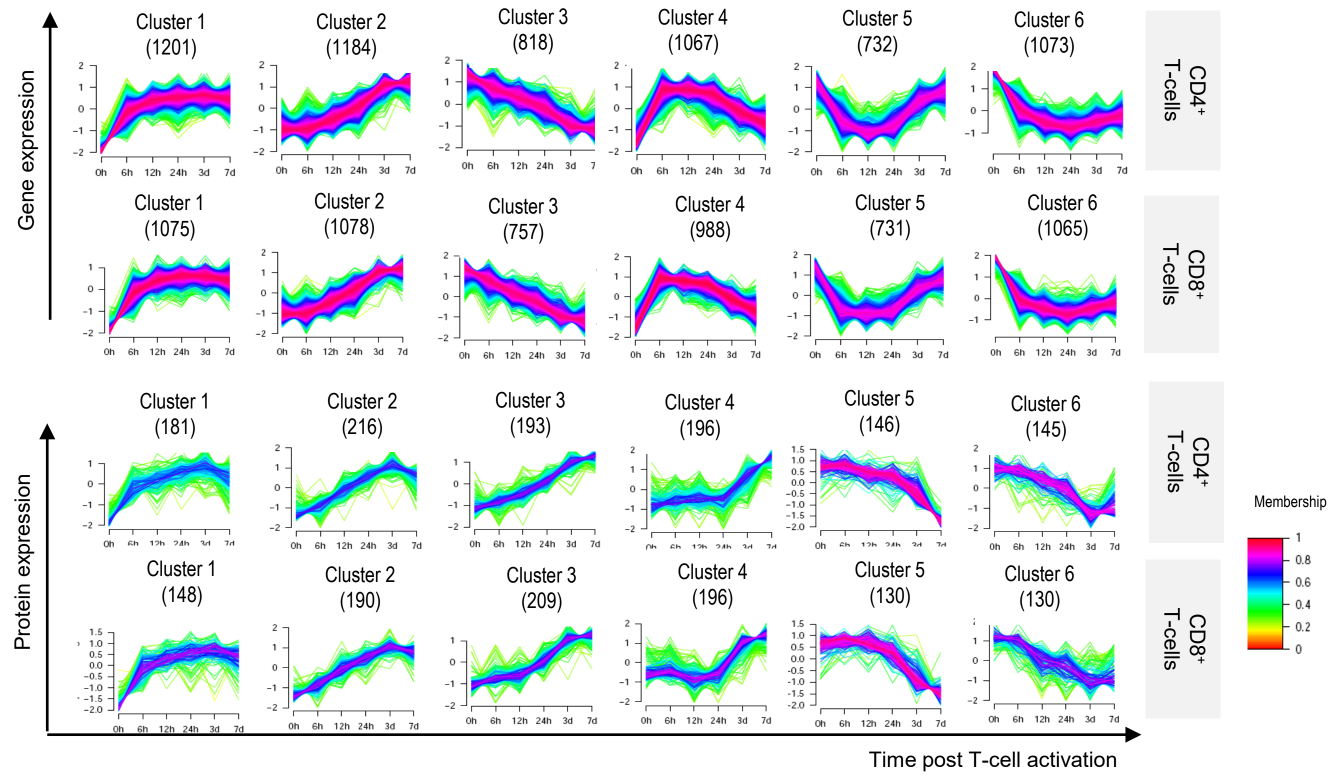


Expression of proteins and mRNA transcripts in each cluster is shown. mFuzz soft clustering was used to identify the co-expression clusters. Each line represent a mRNA/protein and the color indicates the membership of each mRNA/protein to the cluster.

### Supplementary Figure 4

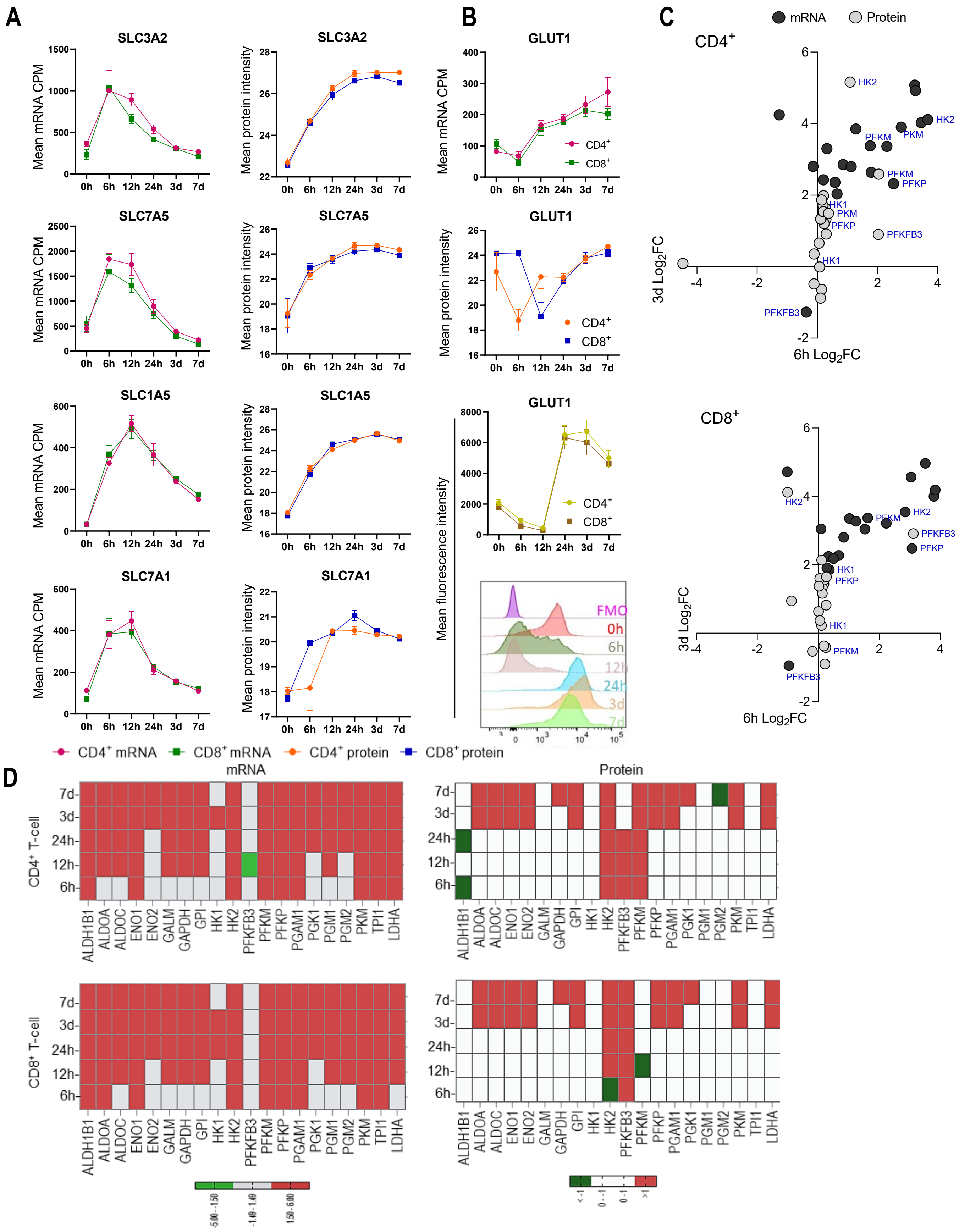

**A.** Expression kinetics of the amino acid transporters SLC3A2, SLC3A5, SLC1A5 and SLC7A1 at both mRNA and protein levels during $CD4^+$ and $CD8^+$ T-cell activation. Expression values of each time point are given as mean and the standard mean of error (SEM). .

**B.** Expression kinetics of the glucose transporter 1 (GLUT1) over the course of $CD4^+$ and $CD8^+$ T-cell activation. Data show values obtained from transcriptomics (mRNA CPM) and proteomics (protein intensity). In parallel, MFI (mean fluorescence intensity) values for cell surface GLUT1 were generated based on flow cytometric analysis and are shown alongside representative histograms of GLUT1 expression on $CD4^+$ T cells.

**C.** Expression of mRNA transcripts and proteins in the glycolytic pathway. Data are given for both $CD4^+$ and $CD8^+$ T-cells and represent enzymes and allosteric regulators regulated at 6 hours (y-axis) and/or 3d (x-axis) post-activation.

**D.** Heatmap representing the expression pattern of mRNA transcripts and proteins in the glycolytic pathway over the entire time course of $CD4^+$ and $CD8^+$ T-cell activation.
